# A qMRI approach for mapping microscopic water populations and tissue relaxivity in the *in vivo* human brain

**DOI:** 10.1101/2024.08.09.606771

**Authors:** Filo Shir, Lee Cohen, Gilad Yahalom, Aviv A. Mezer

## Abstract

Quantitative magnetic resonance imaging (qMRI) enables non-invasive mapping of brain tissue microstructure and is widely used for monitoring various physiological and pathological brain processes. Here, we introduce a qMRI approach for enriching the microstructural characterization of the sub-voxel environment. Inspired by pioneering magnetization transfer (MT) models, this approach employs MT saturation to differentiate between various water populations within each voxel. Our *in vivo* results align well with theoretical predictions and are reproducible using standard qMRI protocols. We present an array of new quantitative maps, highlighting different aspects of the tissue’s water. Furthermore, by manipulating the effective water content and relaxation rate with MT, we approximate within the voxel the tissue relaxivity. This property reflects the dependency of R1 on the macromolecular tissue volume (MTV) and is associated with the lipid and macromolecular composition of the brain. Our approach also enables biophysically-informed modulation of the R1 contrast, resulting in a set of unique cortical profiles. Finally, we demonstrate the effectiveness of our technique in imaging the common pathology of white matter hyperintensities (WMH), revealing tissue degradation and molecular alterations.

## Introduction

Quantitative Magnetic Resonance Imaging (qMRI) techniques have revolutionized our ability to probe the microstructure of brain tissue non-invasively^1,2^. Different qMRI parameters can be estimated, each with distinct biophysical interpretation. Among these techniques, the observed longitudinal relaxation rate (R1) is extensively utilized, enabling the monitoring of a range of physiological and pathological brain processes^1,2^. R1 was shown to be sensitive to different aspects of brain tissue microstructure such as myelination, iron content and water content^3–6^. In addition, qMRI’s ability to assess the tissue water content (WC) presents a distinct advantage in detecting developmental, aging, inflammatory, and edematous changes in the brain^7–13^. The complement parameter, the non-water content (1-WC), represents the macromolecular tissue volume (MTV), i.e. the volume fraction of brain tissue^9^. The dependencies between qMRI parameters were shown to reveal additional information on brain tissue microstructure^14^. Particularly the tissue relaxivity, defined as the influence of changes in the tissue fraction on relaxation, can be estimated through the linear dependency of the observed R1 on MTV across voxels. Tissue relaxivity was shown to be sensitive to the membrane lipid composition of the tissue and to reveal molecular alterations associated with aging ^14^.

The R1, WC, MTV and tissue relaxivity estimation approaches, while invaluable, operate under the simplifying assumption of homogeneity, overlooking the heterogeneous nature of brain tissue^15–19^. This assumption implies modeling a single-exponential relaxation process of a single water pool, resulting in single R1, MTV, and WC values per voxel. This makes it impossible to measure the tissue relaxivity, i.e. the R1-MTV dependency, within the voxel, as at least two data points are required for fitting a line. Currently, linear models to analyze the tissue relaxivity incorporate estimates from all voxels within a specific region of interest (ROI), resulting is a single estimation of tissue relaxivity per ROI^14^. In contrast to the assumption of homogeneity, brain tissue’s sub-voxel makeup exhibits a heterogeneous nature, of multiple water environments with distinct relaxation properties^15–19^. Most cellular water is relatively free, but some is bound and exchanges energy with macromolecular protons. More elaborated qMRI approaches, such as quantitative magnetization transfer (qMT) techniques^20–28^, consider the multi-exponential relaxation of these diverse water environments. However, these models have not been employed in the context of tissue relaxivity and water content mapping. This limits the ability to infer valuable insights into tissue organization at the voxel-wise level.

Assessing local tissue organization at a voxel-wise level based on relaxivity and water environments is crucial when investigating different pathologies. Analyzing tissue relaxivity across the entire white matter, for instance, could mask the presence of white matter hyperintensities (WMH). These are commonly observed in the aging brain and appear as bright lesions in fluid attenuated inversion recovery (FLAIR) imaging^29–31^. WMH are associated with cognitive decline, triple the risk of stroke, and double the risk of dementia^31^. The underlying pathology of WMH includes vascular degeneration, gliosis, infarcts, and microstrokes^30^. Insights into the physiological origins of WMH largely come from post-mortem studies, posing challenges in analyzing water content changes in this pathology. Neuroimaging studies, mainly based on diffusion MRI, suggest that water content changes represent the early and more reversible stages of WMH^29^. Complementing the non-invasive imaging of pathologies like WMH with measurements of tissue relaxivity and water environments at the single voxel level could enhance our understanding of the pathological microstructural heterogeneities and their biophysical underpinnings.

Addressing these challenges, we introduce a novel qMRI approach designed to directly image microscopic water populations and voxel-wise tissue relaxivity within the *in vivo* human brain. This approach utilizes Magnetization transfer (MT) saturation to eliminate the contribution of some of the water protons to the MRI signal. We complement this acquisition with a simple single-exponential biophysical model, influenced by pioneering MT models^32–34^. This model allows to counterpart the standard observed R1, MTV and WC estimations with additional R1sat, MTVsat and WCsat estimations per voxel, acquired under the influence of MT saturation. While there exist more advanced MT models^20–26,35^, we find this model useful for estimating new biophysical parameters which combine several qMRI measurements. It provides access to different water environments within the imaging voxel, based on straightforward data acquisition and parameter fitting approaches. In addition, the MT saturation alters the effective tissue fraction of the voxel, and thereby also the observed R1^32–34^. Consequently, this methodology can be used to estimate the tissue relaxivity within the imaging voxel, offering a tissue relaxivity map for the first time.

In this paper, we present the theoretical framework, technical implementation and empirical validation of our methodology. We also propose different approaches for accelerating its acquisition. By leveraging the voxel-wise mapping of the tissue relaxivity, we generate different biophysically-informed R1 contrasts with distinct cortical profiles. Finally, we demonstrate the applicability of our approach for biophysical characterization of white matter hyperintensities. Therefore, our non-invasive approach offers new insights into the microscopic heterogeneity of brain tissue.

## Theoretical background

### Two-site exchange model

For the biophysical modeling of our approach, we revisited the pioneering magnetization transfer (MT) model of Forsén and Hoffman, presenting an exchange system involving two sites A and B^33^:

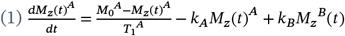

is the longitudinal magnetization, and are the exchange rates and is the steady-state free protons magnetization without irradiation.

Under the assumption of complete saturation of the macromolecular spins (*M*_*z*_^*B*^(*t*) = 0) and no direct effect on the liquid spins ^33^:

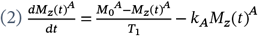

While these assumptions can lead to oversimplification of the model, we will test their usefulness for characterizing the influence of MT on R1 and water content mapping (see Results section for further investigation of these assumptions).

If the saturation of B can be assumed to occur instantaneously at the time t=0, when, the solution for the longitudinal magnetization (we omit here the A notation) ^33^:

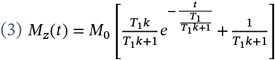

The system obtains a new equilibrium value, with less available equilibrium magnetization^36^:

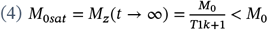

This new equilibrium value is attained through an exponential decay with the time constant *T*1_*sat*_:

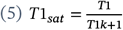

Therefore, this saturated system has higher observed R1 (*R*1_*sat*_= 1/*T*1_*sat*_) compared to the standard observed R1 (1 = 1/*T*1):

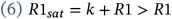

Combining eq. 4 and 5 it can be shown that^32^:

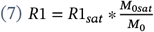

### Mapping and based on the variable flip angle technique

The variable flip angle technique allows to measure *R*1 and *M*_0_(and consequently and, see sub-section “Calibrating M0 for measuring WC and MTV”)^37–39^. This technique involves fitting at least two spoiled gradient echo (SPGR) scans acquired with different flip angles to the Ernst equation^40^:

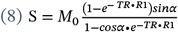

Where TR is the repetition time and α is the effective flip angle following B1+ correction.

We hypothesized that by modifying the standard variable flip-angle (VFA) technique and incorporating MT irradiation for each SPGR scan, the observed and can be measured (and consequently and, see sub-section “Calibrating M0sat for measuring WC and MTV under the influence of MT”). Under the assumption of a single-exponential observed R1, the SPGR signal equation with MT can be described by a modification of the Ernst equation^32,41^:

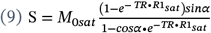

Where and represent and under the influence of MT. Therefore, we hypothesized that by acquiring at least two MT-SPGR scans with different flip angles, and can be evaluated.

### Mapping and based on a fast implementation

Here we will show that instead of using several MT-SPGR scans and the VFA approach, a fast implementation of *M*_0*sat*_ and *R*1_*sat*_ mapping can be achieved with a single MT-SPGR scan (and consequently and mapping, see sub-section “Calibrating M0sat for measuring WC and MTV under the influence of MT”). This fast implementation requires, and bias fields maps estimated independently. It is based on eq. 9 in which there are two unknowns (M0sat and R1sat). Under the constraint of eq. 7, that allows to express as function of *R*1_*sat*_, *R*1and *M*_0_, a single unknown parameter is left:

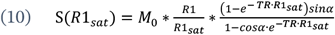

Where S is the signal, TR is the repetition time and α is the effective flip angle following B1+ correction. and are estimated independently from SPGR scans (eq. 8). Eq. 10 allows for the estimation of *R*1_*sat*_. *M*_0*sat*_ can then be calculated based on eq. 7. Therefore, *M*_0*sat*_ and *R*1_*sat*_ can be estimated based on a single MT-SPGR scan.

### Calibrating M0 for measuring WC and MTV

The proton density (PD) can be calculated based the equilibrium magnetization (*M*_0_) after accounting for the receive-coil sensitivity (B1-)^8^:

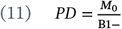

Assuming the MRI signal is generated by water protons, and that brain ventricles contain 100% water, PD values across the brain can be scaled relative to the median PD value in the ventricles (*PD*_*ventricles*_) for producing apparent water content (WC) estimates^9^:

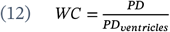

The non-water content represents the macromolecular tissue volume (MTV):

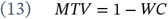

### Calibrating for measuring WC and MTV under the influence of MT

Estimating PD under MT saturation 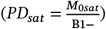, there should be less available longitudinal magnetization (eq. 4) and therefore suppression of some of the signal-producing protons. This should result in lower values, and consequently lower apparent water content (*WC*_*sat*_):

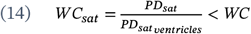

And substituting the WC definition based on M0 into eq. 7:

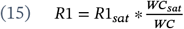

With less signal-producing water protons following MT saturation, the observed non-water tissue fraction relative to water is expected to effectively increase:

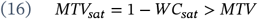

### Mapping different water populations under the influence of MT

We postulated that two different water populations can be estimated in each voxel. We defined the saturated water content (SWC) as the water eliminated from the signal following MT saturation:

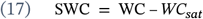

Whereas the unsaturated water content (USWC) is the water that continues to contribute to the signal in the presence of MT:

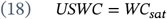

The saturated water fraction can be defined as the fraction of saturated water to total water:

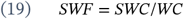

### Voxel-wise estimations of the tissue relaxivity and the conditional R1 distributions

The tissue relaxivity was previously defined as the linear dependency of R1 on MTV across voxels^14^:

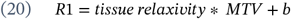

Where the tissue relaxivity is the slope 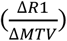 and b is the intercept.

We hypothesized that the tissue relaxivity can be estimated per voxel based on the relative change in the observed R1 and MTV following MT:

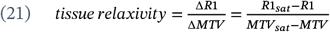

We further defined the conditional distributions of R1 as the expected R1 values based on the linear model in eq. 20:

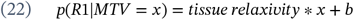

Where b can be calculated as:

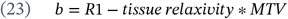

## Methods

### Human Subjects

The study includes 7 healthy young adults (ages 24-33 years, 2 males). For the WMH analysis, additional 5 healthy older adults (ages 74-79 years, 2 males) were selected from an initial sample of 27 subjects (see Methods section “WMH analysis” for selection criteria). Volunteers were recruited from the community surrounding the Hebrew University of Jerusalem. The experimental procedure was approved by the Helsinki Ethics Committee of Hadassah Hospital, Jerusalem, Israel. Written informed consent was obtained from each participant prior to the procedure.

### MRI acquisition

Data were collected on a 3T Siemens MAGNETOM Skyra scanner equipped with a 32-channel head receive-only coil at the ELSC Neuroimaging Unit at the Hebrew University. One of the young subjects was scanned twice for the entire protocol.

SPGR scans (TE/TR=3.34/19 ms) were acquired with variable flip angles (α = 4°, 10°, 20°, and 30°). The scans resolution was 1 mm isotropic (for all subjects except one, see below). For the TR sensitivity analysis, one young subject underwent supplementary scans with a TR of 27 ms.

MT-weighted SPGR scans (TE/TR = 3.34/ 27 ms, α = 10°) were acquired for all subjects. For young subjects, additional variable flip angle MT-weighted SPGR scans were acquired (α = 4°, 20°, 30°, TE/TR = 3.34/ 27 ms, 1mm isotropic). The same MT pulse was used in all these scans (gaussian pulse, 10 ms, 500°, 1.2 kHz, off/on resonance amplitudes ratio=2.034). The scans resolution was 1 mm isotropic (for all subjects except one, see below).

For one of the young subjects, the SPGR and MT-SPGR scans were acquired in a 1.6 mm isotropic resolution. For this subject 4 additional MT-weighted SPGR scans were acquired with different MT pulse amplitudes (off/on resonance amplitudes ratio ranging from 1.9-2.64). TE/TR in all MT-weighted scans for this subject was 3.34/34 ms, and α = 10°.

For two of the young subjects, additional variable flip angle MT-weighted SPGR scans were acquired with higher MT pulse amplitude (for one subject: off/on resonance amplitudes ratio=2.7, TE/TR=3.34/36 ms, α = 10°, for the second subject: off/on resonance amplitudes ratio=2.25 and 2.445, TR=28 and 32 ms correspondingly, TE=3.34 ms, α = 10°).

For B1+ mapping, we acquired spin-echo inversion recovery scan with an echo-planar imaging read-out (SEIR-epi) for all subjects. This scan was done with a slab-inversion pulse and spatial-spectral fat suppression. For SEIR-epi, the TE/TR were 49/2,920 ms. The TIs were 200, 400, 1200, and 2400 ms. We used 2-mm in-plane resolution with a slice thickness of 3 mm. The EPI readout was performed using 2× acceleration.

For segmentation, 3D magnetization-prepared rapid gradient echo (MPRAGE) scans were acquired (1 mm isotropic, TE/TR=2.98/2300 ms).

For WMH segmentation we acquired 3D T2-Fluid Attenuated Inversion Recovery (FLAIR) with TR /TE/TI= 9000/ 72/2500 ms, flip angle = 150° and 0.75 × 0.75 × 3.6 mm resolution.

### Evaluation of qMRI parameters

Whole-brain WC, MTV and R1 mapping, together with field maps of excitation bias (B1+) and receive-coil bias (B1-), were computed using the mrQ software^9^ based on SPGR and SEIR scans (see eq. 8).

MT-SPGR scans were registered to the imaging space of WC, MTV and R1 using a rigid-body alignment.

Whole-brain WCsat, MTVsat and R1sat mapping were computed based on variable flip angle MT-SPGR scans (α = 4°, 10°, 20°, and 30°) with a modified version of mrQ (see eq. 9). This modified version uses the bias maps (B1- and B1+) and the CSF mask calculated for WC, MTV and R1 (based on the SEIR and SPGR scans).

We also calculated WCsat, MTVsat and R1sat with the fast implementation (eq. 10), based on a single MT-SPGR scan (α = 10°). This implementation requires the estimations of M0, R1, bias maps (B1- and B1+) and the CSF mask obtained from the SEIR and SPGR scans.

MTR maps were produced based on the equation: MTR=(S_syn_-S_MT_)/S_syn_, where S_MT_ is the MT-SPGR scan (α = 10°) and S_syn_ is a synthetic SPGR scan with the same TR and flip angle as S_MT_ (computed from WC, R1 and the bias maps).

Segmentation: Whole-brain segmentation was computed automatically using the FreeSurfer segmentation algorithm (v6.0)^42^. MPRAGE scan registered to the R1 space using a rigid-body alignment was used as a reference. Sampling of 9 equidistance points along the cortex (−0.4-1.2) was done with FreeSurfer’s function mri_surf2vol^42^.

### Direct effect calibration

The theoretical model was developed under the assumption of no direct effect (DE) on the liquid spins (eq. 2). Notably, this assumption is usually imprecise for *in vivo* data.

In the absence of a direct effect, there should be no observed MT saturation in the ventricles, where there is approximately 100% water. Therefore, there should be similar water content estimations with and without MT in the ventricles (*WC*(*ventricles*) = *WC*_*sat*_(*ventricles*)) ^32,33^. Hence, the magnitude of the direct effect can be estimated as follow:

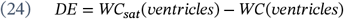

To account for the direct effect, DE was estimated based on eq. 24, by taking the median WC and WCsat values in the CSF mask. Voxels in which WCsat values were higher than 1 were excluded from the CSF mask.

The uncalibrated WCsat maps 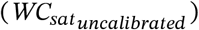 were then calibrated to have the same median value as WC in the ventricles, by subtracting DE for all voxels in the whole-brain map:

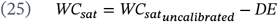

For calibration of MTVsat we used the calibrated WCsat (MTVsat=1-WCsat). This calibration was performed for WCsat and MTVsat maps obtained either by the full fitting in eq. 9 or by the fast implementation in eq. 10.

### Evaluation of biophysical parameters

We calculated different biophysical parameters based on R1, MTV, R1sat and MTVsat. Whole-brain tissue relaxivity maps were produced based on eq. 21. Maps of different water populations were generated based on WC and WCsat following eq. 17-19. The conditional distributions of R1 were calculated with eq. 22-23. Most of the presented analyses were based on variable flip angle mapping (eq. 8-9). Cases in which the fast implementation of our approach was employed (eq. 10) are explicitly indicated.

ROI-based tissue relaxivity estimates were computed as described previously^14^. For each ROI, we extracted the MTV values from all voxels and pooled them into 36 bins spaced equally between 0.05 and 0.40. This was done so that the linear fit would not be heavily affected by the density of the voxels in different MTV values. We removed any bins in which the number of voxels was smaller than 4% of the total voxel count in the ROI. The median MTV of each bin was computed, along with the median of the qMRI parameter. We used these data points to fit the linear model across bins using eq. 20. The slope of the line is the whole-ROI tissue relaxivity.

### WMH analysis

We evaluated an initial sample of 27 subjects aged over 45 for whom the protocol for producing the fast implementation of our approach was successfully acquired (SPGR scans, SEIR scans and a single MT-SPGR scan) together with FLAIR and MPRAGE scans. WMH segmentation was done manually based on the FLAIR scan. Normal appearing white-matter (NAWM) was segmented with FreeSurfer ^42^ using the MPRAGE scan as a reference. These segmentations were registered to the qMRI imaging space with a rigid-body alignment. To identify subjects with WMH pathology, selection criteria included a lesion load (WMH/NAWM) greater than 0.02 and higher FLAIR values in WMH compared to NAWM, indicating hyperintensity. This selection criteria resulted in 5 subjects which are included in the final analysis. Parameters were estimated with the fast implementation of our approach (eq. 10).

## Results

### New MRI contrasts agree with the model’s prediction

Initially, we examined the effect of MT-weighting on the observed longitudinal relaxation, tissue fraction, and water content. For this aim, we applied the variable flip-angle technique for MT-SPGR scans (α = 4°, 10°, 20°, and 30°) and fitted a single-exponential model (eq. 9). This allowed for the estimation of R1sat (R1 with MT-weighting), MTVsat (MTV with MT-weighting), and WCsat (WC with MT-weighting) in the *in vivo* human brain. Figure 1 shows an example of these new qMRI parameters in a single subject, compared to the standard non MT-weighted parameters (Fig. 1ab). Notably, our findings highlight a significant change in the acquired contrasts due to the MT-weighting. R1sat, MTVsat, and WCsat exhibit unique distributions within the brain, distinct from those of R1, MTV, and WC respectively (Fig. 1c, p<10^−4^ and averaged test statistics>0.6 for the two-sample Kolmogorov–Smirnov test comparing the distributions across all brain voxels in n=6 subjects). These results suggest the potential of the new contrasts to unveil unique microstructural information.

**Figure 1:**
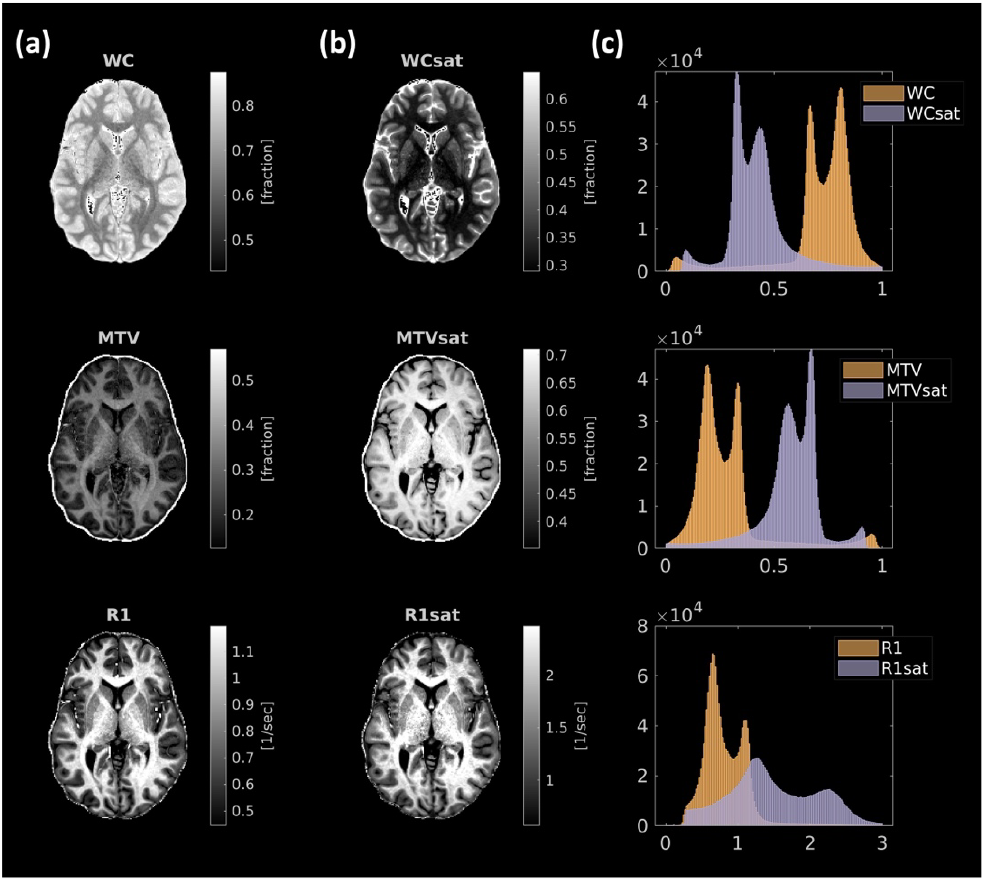
The effect of MT-weighting on WC, MTV and R1 mapping. **(a)** The observed WC, MTV and R1 contrasts are presented for a representative human subject. **(b)** The effect of MT on these observed contrasts is shown for the same human subject: WCsat (WC with MT), MTVsat (MTV with MT) and R1sat (R1 with MT). **(c)** Comparison between these contrasts is shown by the distribution of values in the entire brain of a single subject. The MT-weighting results in the generation of novel and distinct qMRI contrasts.

We further validated that the contrast changes due to the MT pulse are not caused by biases in the acquisition and estimation. First, we assessed the goodness of fit for a single-exponential model applied to MT-SPGR data, which may underlie a multi-exponential relaxation process. We compared the model performances on MT-SPGR data to its performances on non-MT SPGR data, for which a single-exponential model if often applied (Fig. 2a). The mean absolute percentage errors (MAPE) were similar when applying the model for both MT-SPGR and non-MT-SPGR scans, indicating that the single-exponential model consistently explained the data (Fig 2b, median MAPE with MT=1.60%, without MT=2.14%, across all voxels in the brains of 6 subjects). Second, MT-weighted scans had to be acquired with a longer TR compared to non-MT-weighted scans. We confirmed that the distinct contrasts observed for R1sat, MTVsat, and WCsat in comparison to R1, MTV, and WC are attributed to the MT pulse rather than the TR difference. We acquired an additional set of MTV, WC and R1 maps with a longer TR, similar to the one used in the MT-SPGR scans. We found that R1 and MTV (1-WC) values remained relatively stable across TR, and were distinct from MTVsat and R1sat values even when all scans were acquired using the same TR (Sup. Fig. 1a). Therefore, the change in contrast following MT does not originate from the TR difference. Lastly, we tested the reproducibility of the new parameters through a scan-rescan experiment (Sup. Fig. 1b). We found consistent parameter estimates between scans, supporting that the unique contrasts of R1sat, MTVsat, and WCsat reflect biophysical information rather than measurement errors.

**Figure 2:**
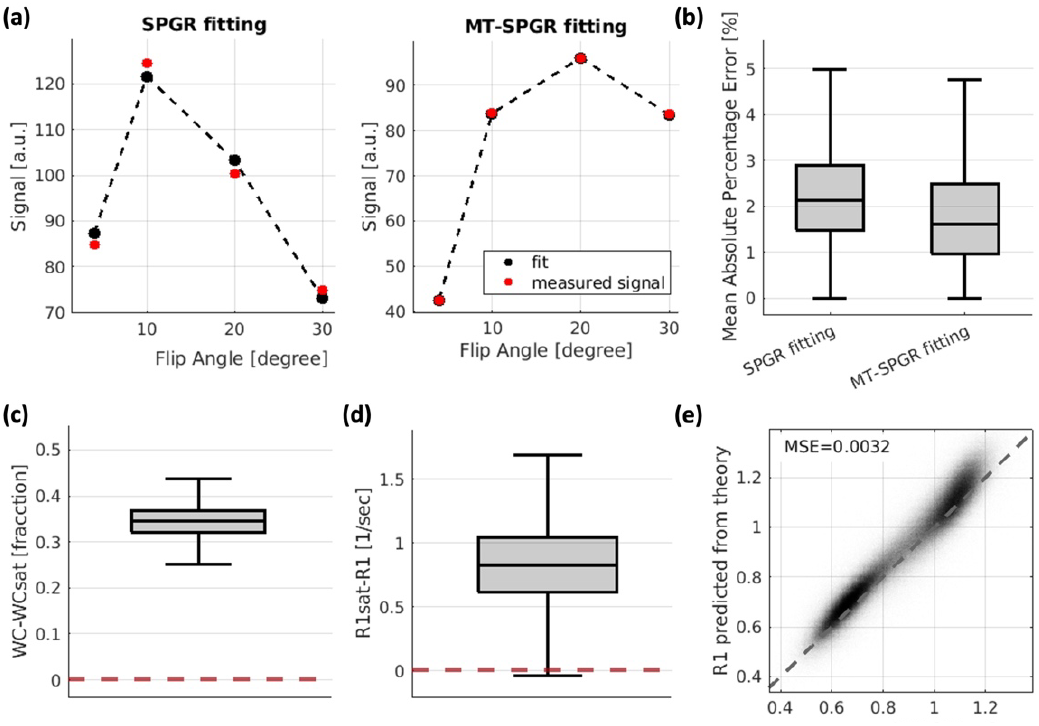
Validating the theoretical basis for the MT-weighted WC and R1 mapping. **(a)** Demonstration of the fitted model with and without MT: the measured signal (black) as function of the flip angle in a single representative voxel in the WM for SPGR scans (right) and MT-SPGR scans (left). The fitted model is in red (eq. 8 and 9 correspondingly) **(b)** Mean Absolute Percentage Error (MAPE) for fitting the measured SPGR and MT-SPGR scans (4 flip angles) to the model of the signal equation (eq. 8 and 9 correspondingly). Results are shown for all voxels in the brains of 6 healthy young subjects. The 25th, 50th, and 75th percentiles and extreme data points are shown for each box. Error levels are similar for both MT and non-MT models. **(c)** The observed water content estimation with MT (WCsat) is lower than the observed water content estimated in the absence of MT (WC). The positive difference in WC-WCsat estimations is shown for all voxels in the brains of 6 healthy young subjects. The 25th, 50th, and 75th percentiles and extreme data points are presented. **(d)** The observed R1 estimated with MT (R1sat) is higher than the observed R1 estimated in the absence of MT (R1). The positive difference in R1-R1sat estimations is shown for all voxels in the brains of 6 healthy young subjects. The 25th, 50th, and 75th percentiles and extreme data points are presented. **(e)** The measured R1 values (x-axis) against the predicted R1 (y-axis) based on the two-site exchange model (eq. 15), including calibration for the direct effect. Results are for all voxels in the brains of 6 healthy young subjects. MSE= mean squared error.

Subsequently, we assessed the agreement of the measured parameters with the theory. In line with the prediction of the two-site exchange model (eq. 14)^32–34^, the measured WCsat is smaller than the total WC, implying the suppression of some water from the signal (Fig. 2c). In addition, consistent with theoretical expectations (eq. 6), R1sat is higher than R1 (Fig. 2d). Furthermore, we identified a strong concordance between the theoretical prediction for R1 based on WCsat, WC, and R1sat (eq. 15) and the R1 measured *in vivo* (MSE=0.0032, across all voxels in the brains of 6 subjects, Fig. 2e). Therefore, the measured R1sat, MTVsat, and WCsat in the *in vivo* brain align with the theoretical expectations, highlighting the relevance of the original MT model to the measured data (the theoretical assumptions regarding partial saturation and the direct effect are tested in Results section “Testing the model’s saturation assumptions”).

### Water population mapping

By combining R1sat, MTVsat and WCsat with the standard R1, MTV and WC, we could generate different biophysical parameters describing the relationships between qMRI measurements. First, the proposed method for mapping WC following MT saturation allowed for the delineation of two water populations within the imaging voxel (eq. 17-19). We estimated the saturated water content as the water eliminated from the signal following MT saturation (SWC = WC - WCsat), and the unsaturated water content (USWC = WCsat) as the water that continues to contribute to the signal in the presence of MT. Additionally, the saturated water fraction can be estimated as the fraction of saturated water to total water (SWF = SWC/WC). The qMRI maps of the saturated and unsaturated water, along with the saturated water fraction (Fig. 3) demonstrate that our approach enriches the description of the sub-voxel water environment.

**Figure 3:**
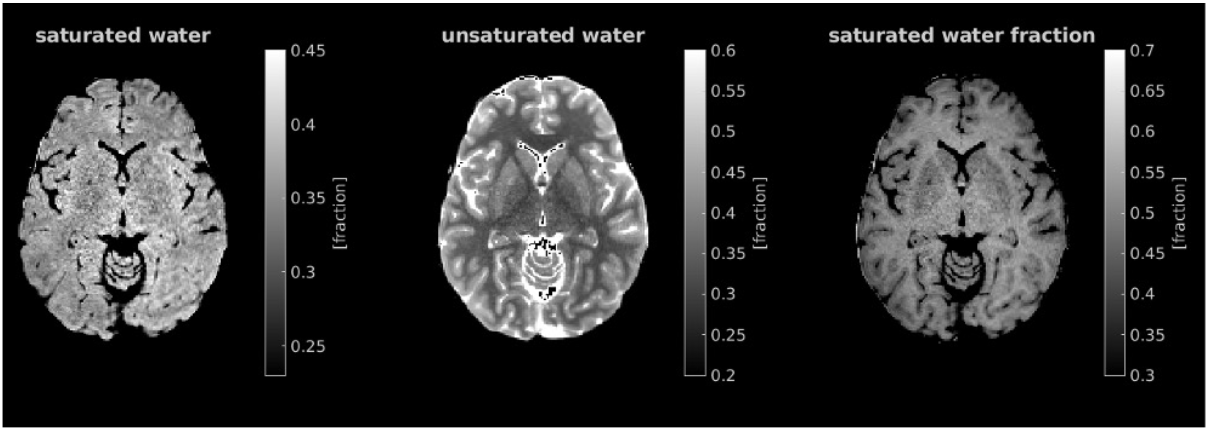
In vivo qMRI of different water populations in the human brain. Whole brain in vivo mapping of the saturated water (left) and the unsaturated water (middle) in a representative subject. The saturated water fraction (SWF), reflecting the ratio of saturated to total water content, is also shown (right).

### Voxel-wise relaxivity

Next, we evaluated the applicability of our approach for voxel-wise tissue relaxivity mapping. The tissue relaxivity represents the sensitivity of relaxation to changes in the tissue fraction, estimated by MTV. Current tissue relaxivity measurements rely on variations in tissue fraction across voxels, thereby limiting the resolution of this measurement to whole ROIs^14^. The MT water saturation changes the observed tissue fraction within voxels, as evident by the increase in MTV following MT (i.e. MTVsat is higher than MTV, Fig. 1). Measuring two observed MTV and R1 values within each voxel, with and without MT, allows to approximate the dependency of R1 on MTV (i.e., the tissue relaxivity) at a voxel-wise resolution (Fig. 4a). We leverage this observation to quantify a map of the local tissue relaxivity (Fig. 4b).

**Figure 4:**
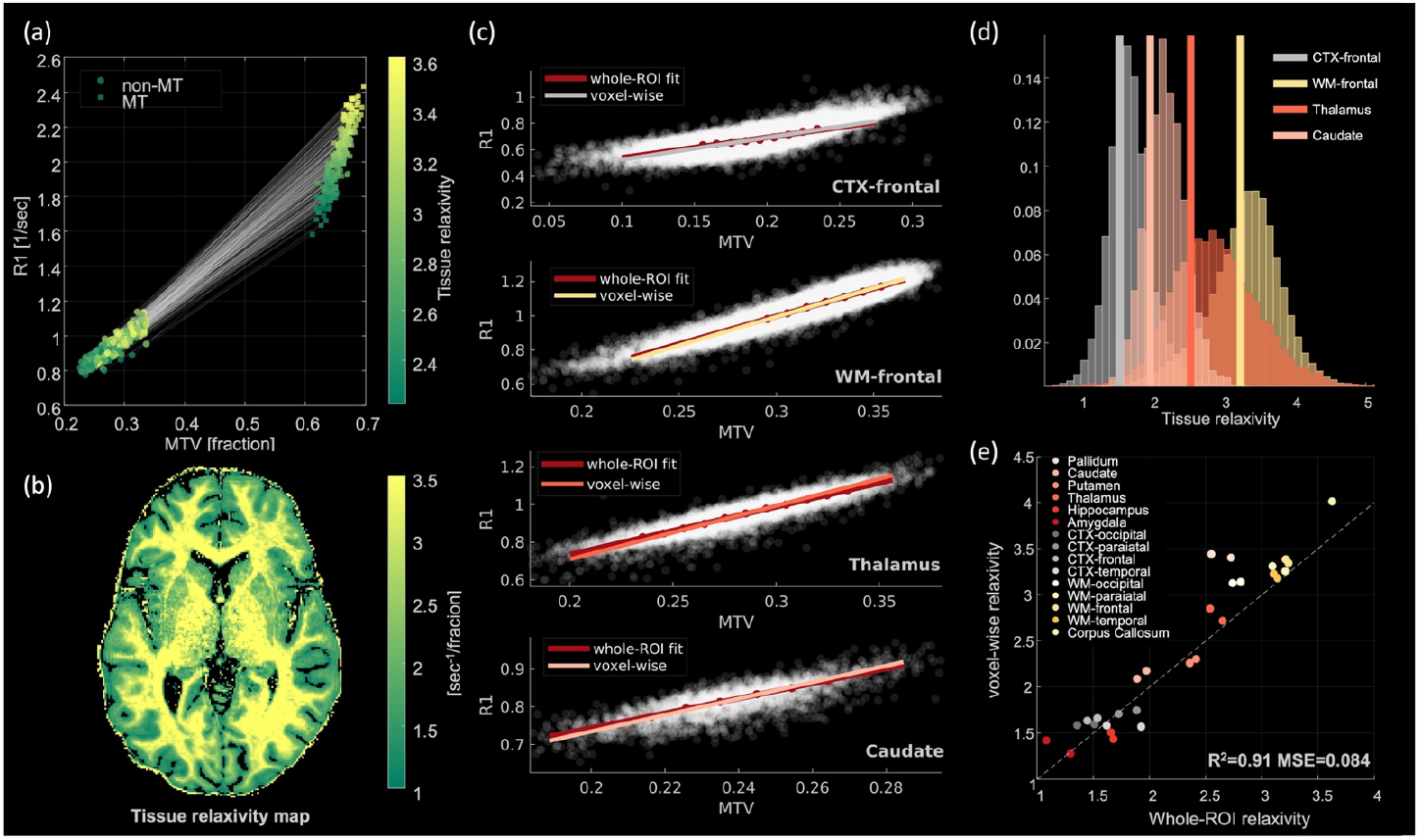
The voxel-wise qMRI map of the tissue relaxivity and its agreement with the whole-ROI approach. **(a)** Estimating the tissue relaxivity (the linear dependency of R1 on MTV) within the voxel; MTV vs. R1 (circles) and MTVsat vs. R1sat (squares) for 250 thalamic voxels. The slope between these two data points is the tissue relaxivity. It is shown by the gray lines, each connect between the saturated and non-saturated measurements of the same voxel. Data points are color-coded by the tissue relaxivity value of each voxel. **(b)** Voxel-wise map of the in vivo tissue relaxivity in a representative subject. **(c)** Comparison of the whole-ROI and the voxel-wise tissue relaxivity for four ROIs in the brain of a representative subject. For each ROI, gray data points show the R1 and MTV values across voxels. Red line represents the whole-ROI tissue relaxivity; MTV values across voxels were pooled into bins, and a linear fit was calculated based on the bins’ median R1 and MTV values (red dots). Whole-ROI tissue relaxivity is the slope extracted from the fit (red line). For each ROI, the voxel-wise tissue relaxivity averaged over all voxels of the ROI is shown on top of the whole-ROI estimation (colored lines). **(d)** Histograms show the distribution of voxel-wise tissue relaxivity values within all voxels of each ROI (different colors). These distributions are centered around the whole-ROI estimation of tissue relaxivity (shown by the vertical lines), indicating the similarity between the whole-ROI and the averaged voxel-wise estimations. **(e)** The significant correlation between the averaged voxel-wise tissue relaxivity and the whole-ROI relaxivity across 15 brain regions in a single subject (for more subjects see Sup. Fig 2). MSE=mean squared error.

Importantly, we find great agreement between the estimations of tissue relaxivity obtained on the whole-ROI and voxel-wise levels. In Figure 4c, the whole-ROI tissue relaxivity is presented for various brain regions, alongside the voxel-wise tissue relaxivity estimation averaged across voxels. The observed dependency of R1 on MTV is consistent for both cases. Figure 4d shows the distribution of voxel-wise tissue relaxivity values within different brain regions. These histograms are centered around the whole-ROI tissue relaxivity estimation. The agreement between the voxel-wise and the whole-ROI tissue relaxivity dependency of R1 on MTV is estimations is also replicated across other brain regions (Fig. 4e, R^2^=0.91), and across subjects (Sup. Fig. 2). These findings indicate that whether assessing changes in R1 and MTV across voxels with the ROI approach or within voxels with MT, we are estimating similar property of relaxivity. Thus, the voxel-wise relaxivity approach provides a good approximation of the previously defined R1-MTV dependency and greatly improves the resolution of this measurement.

### Fast implementation of the new approach

The acquisition protocol of our new technique requires two sets of SPGR scans with variable flip angles, with and without MT pulse. While our results are shown for sets of four flip angles (4°,10°,20°,30°), we show that sets of two flip angles (4°,20°) are sufficient for accurate parameter estimation (Sup. Fig. 3). Standard qMRI protocols often contain variable flip angle SPGR scans with at least two flip angles^43,44^. However, these protocols usually include only a single MT-weighted SPGR scan^35,45^. We propose an adaptation of our technique to accelerate its acquisition and make it applicable for such standard qMRI protocols (see “Mapping and based on a fast implementation” in Theoretical Background). The acceleration is based on the theoretical relationship between R1, R1sat, M0sat and M0 (eq. 7), as an additional constraint in the fitting process. This relationship was found to describe our *in vivo* results very well (Fig. 2e, in this case we used eq. 15, which directly follows eq. 7). Figure 5 compares between R1sat, MTVsat and the tissue relaxivity estimated with the full protocol, including four variable flip angle SPGR scans with MT pulse, to the same parameters estimated by using equation 10 based on a single MT-weighted SPGR scan. The full protocol and the fast implementation provide very similar parameters estimations (Fig. 5). Moreover, the fast implementation of our approach is reproducible across scans (Sup. Fig. 4).

**Figure 5:**
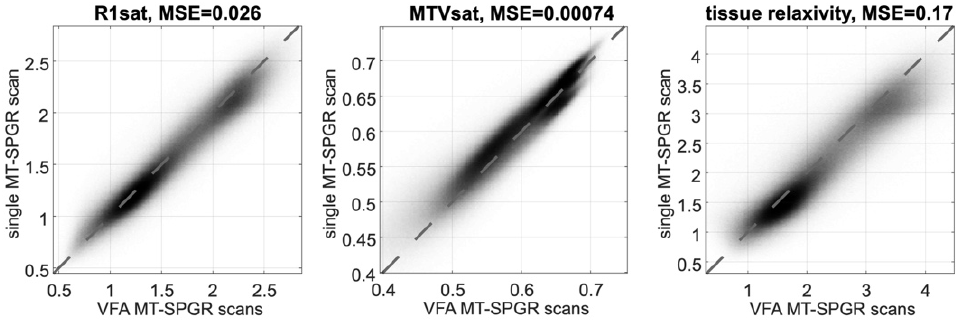
Fast implementation of the tissue relaxivity mapping based on biophysical constraints. R1sat, MTVsat and tissue relaxivity fitted with four variable flip angles (VFA) MT-SPGR scans (x-axis) are well approximated by the fast implementation of our approach, based on a single MT-SPGR scan (y-axis, eq. 10). Values are for all the voxels in the brains of 6 subjects. MSE=mean squared error.

Therefore, by adding only a single MT-weighted SPGR scan to the standard R1 and MTV mapping protocol, both MTVsat and R1sat can be evaluated, together with MTV and R1. This is possible due to the built-in assumption that these properties have a known dependency (eq. 15). Combining the parameters obtained by this approach, allows to estimate the biophysical properties of the tissue relaxivity, the unsaturated and saturated water contents and the saturated water fraction based on a widely used qMRI protocol^43,44^.

### Testing the model’s saturation assumptions

The theoretical formulation we rely on assumes full saturation of the macromolecular spins and does not take into account the direct effect on the liquid spins (see “Two-site exchange model” in Theoretical Background)^32–34^. These conditions are usually not satisfied in standard *in vivo* MRI scans^20,46^. We therefore assessed the effect of these simplifying assumptions on our *in vivo* measurements.

To address the direct saturation of the liquid spins, we estimated the magnitude of this effect based on measurements in the ventricles (see “Direct effect calibration” in Methods). We assumed the CSF approximately does not contain macromolecules, and therefore MT-related changes observed in the ventricles can be attributed to the direct effect (eq. 24). Using this estimation of the direct effect, we calibrated our measurements in the whole brain, by requiring that WC and WCsat will have the same median values in the ventricles (eq. 25). To evaluate the efficiency of the direct effect calibration, we compared between our *in vivo* results and the theoretical model in which we assumed no direct effect. Notably, the agreement between the theoretical R1 model (eq. 15) and the measured R1 improved greatly by the direct effect calibration, as evident by the reduction in mean squared error (MSE=0.019 without calibration and with calibration, Fig. 6a). This finding was replicated across subjects (Fig. 6b).

**Figure 6:**
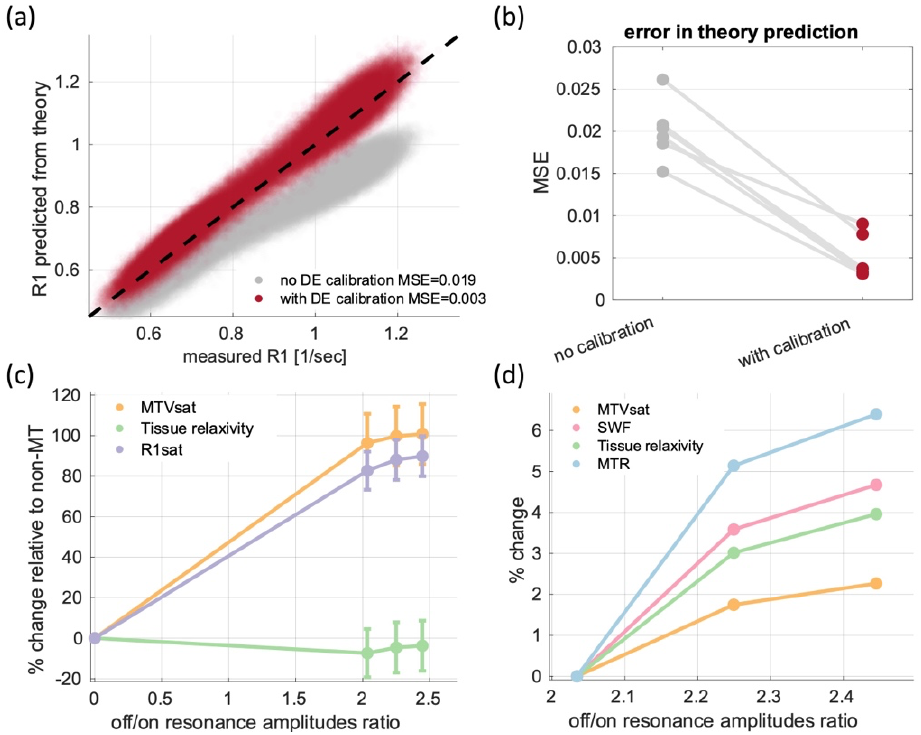
Testing the assumptions of the biophysical model. **(a)** The influence of the direct effect (DE) calibration in a representative subject: the measured R1 (x-axis) is compared against the R1 predicted by the theoretical model (eq. 15, y-axis). This comparison is shown without DE calibration (gray) and after the DE calibration (eq. 25, in red). Values are shown for all voxels in the brain of a single subject. Dashed line is the identity line. R1 prediction by the biophysical model improved when correcting for the direct effect, as evident by the lower mean squared error (MSE). **(b)** The efficiency of DE calibration across subjects is demonstrated by consistently lower MSE values when comparing the measured R1 to the theoretical R1 model with DE calibration, in contrast to the MSE values without calibration (n=6 subjects). **(c)** Comparing parameters estimated using different MT saturation strengths to those estimated without MT: percentage change in R1sat, MTVsat, and tissue relaxivity (y-axis) across various MT pulse amplitudes (off-resonance amplitude to on-resonance amplitude ratio, x-axis). Changes are relative to the estimation of these parameters without MT (R1, MTV, and whole-ROI tissue relaxivity, respectively). Parameters were calculated with the fast implementation of our approach, results are shown in the WM of a single subject (for more subjects see sup. Fig. 5). Error bars indicate standard deviations. Compared to R1sat and MTVsat, the tissue relaxivity estimation with MT is more similar to non-MT estimation. **(d)** The changes in parameters estimations across different MT saturation strengths. percentage change in MTVsat, voxel-wise tissue relaxivity, SWF and MTR (y-axis) for various MT pulse amplitudes (off-resonance amplitude to on-resonance amplitude ratio, x-axis). Changes are relative to the lowest MT pulse amplitude acquired. Parameters were calculated with the fast implementation of our approach, results are shown in the WM of a single subject (for more subjects see sup. Figure 5). All tested parameters exhibit sensitivity to partial saturation effects.

The implications of partial saturation of the macromolecular spins were evaluated by varying the MT pulse strength. Working on conventional Siemens scanner, we could manipulate the strength of the pulse by changing the amplitude of the MT off-resonance pulse relative to the on-resonance pulse. We identified an increase in MTVsat with increasing amplitude of the MT pulse (Fig. 6c, see Sup. Fig. 5a for more subjects). This indicates that as the macromolecular pool becomes more saturated, more water are eliminated from the signal. R1sat and tissue relaxivity also increased with the MT pulse strength, implying for the sensitivity of these parameters to partial saturation effects as well. Relative to the estimations of R1 and MTV without MT, R1sat and MTVsat showed maximal increase of 103% and 115%, correspondingly, with increasing amplitude of the MT pulse (in the WM, Fig. 6c). Importantly, the tissue relaxivity estimation was more stable relative to the non-MT estimation, with maximal change of 6% for all MT amplitudes tested (in the WM, Fig. 6c). We then tested the percental change in parameters estimations between different MT pulse amplitudes. We find that the tissue relaxivity, the SWF and MTVsat increased with the amplitude of the MT pulse, indicating they are affected by the simplifying assumption of complete macromolecular saturation (Fig. 6d, see Sup. Fig. 5b for more subjects). These partial saturation effects were similar in size to the ones observed for MTR.

To conclude, although some of the assumptions in our theoretical model are imprecise, we verified that the direct affect is accounted for. The presence of partial saturation effects calls for careful consideration when comparing estimates acquired with different MT pulse characteristics.

### Biophysically-informed synthetic R1 contrasts

After establishing the reliability and limitations of the voxel-wise tissue relaxivity measurement, we demonstrate its applicability for studying the biophysical sources of the R1 contrast in the brain. The observed R1 is known to be highly sensitive to the water content^14,15,19^. Consequently, a considerable portion of the R1 variability across the adult brain, particularly between cerebrospinal fluid (CSF), gray matter (GM) and white matter (WM), can be attributed to regional differences in the tissue water content. Thus, our objective was to examine the hypothetical scenario of R1 contrast in the absence of these regional water content differences. By leveraging the tissue relaxivity measurement, we generated synthetic R1 contrasts with uniform water content throughout the brain. The tissue relaxivity reflects the linear dependency of R1 on the non-water content, estimated by MTV (eq. 20). Employing this linear equation enables the prediction of R1 for different water content values (eq. 22). Figure 7a shows the synthetic R1 contrasts obtained for various MTV values, held constant throughout the brain.

**Figure 7:**
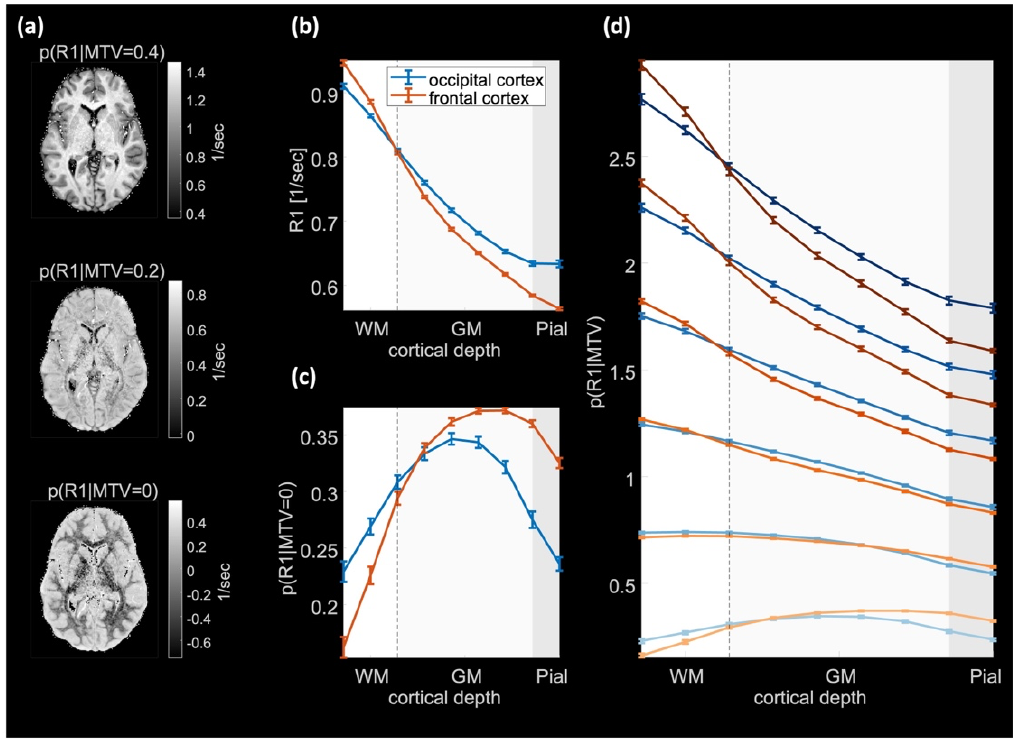
The conditional distributions of R1 and their unique cortical profiles. Demonstration of the conditional R1 distributions p(R1|MTV=x) for three values of MTV (x=0, 0.2 and 0.4) in a representative subject. Each contrast represents the regional changes in R1 when fixing MTV values for the entire brain, and was estimated by the linear relaxivity equation (eq. 22). The contrast inversion between GM and WM is observed. **(b)** The cortical profiles of standard R1 values (y-axis) sampled at 9 equidistance points along the cortex from WM to pial (x-axis), in the occipital cortex (blue) and the frontal cortex (red) **(c)** The cortical profiles of p(R1|MTV=0), indicating the non-MTV variability across the cortex **(d)** The conditional R1 distributions for different values of MTV (y-axis) as function of cortical depths (x-axis). Profiles of the occipital cortex (blue) and the frontal cortex (red) are shown. Darker colors represent increasing MTV values (ranging from 0 to 1). The cortical profiles of R1 vary with the fixed value of MTV. Errorbars in the entire figure show the standard error of the mean (n=6 subjects).

We defined these synthetic contrasts as the conditional distributions of R1 in the brain, for fixed tissue fraction (p(R1|MTV=x), eq. 23). Interestingly, at very low MTV values (indicating high water content), we observe a contrast inversion, with higher R1 in the GM compared to the WM. This is shown in figure 7a for the conditional distribution of R1 when MTV is set to 0, i.e. p(R1|MTV=0). Therefore, there is a larger proportion of residual R1 unexplained by MTV in the GM compared to the WM. By gradually increasing the simulated water content, we generated a series of synthetic R1 contrasts (sup. movie 1). We find that as MTV increases (water content decreases), the contrast between GM and WM is inverted. To further explore the conditional distributions of R1, we evaluated their cortical profiles from deep WM to the pial surface (Fig. 7b-d). The characteristic pattern of R1 cortical profiles exhibits monotonically decreasing values from the WM to the pial surface (Fig. 7b)^3^. We found that different fixed MTV values yield distinct R1 profiles along the cortex. For example, for p(R1|MTV=0), the cortical profile of R1 obtains an inverted U-shape, with higher values in the mid-cortex compared to the WM and pial surface (Fig.7c). In contrast, for higher MTV values the characteristic shape of the cortical profiles is restored (Fig. 7d).

Moreover, we compared between the conditional R1 profiles of the frontal and occipital cortex. We find that the variability in the profiles of these cortical regions depends on the chosen MTV value (Fig. 7c-d). For example, for p(R1|MTV=0), the conditional R1 in the frontal cortex is lower towards the WM and higher towards the pial compared to the occipital cortex. Increasing MTV, the differences in R1 values between these cortical regions diminish, and for p(R1|MTV=1) we find the opposite trend (Fig. 7d). Therefore, the conditional R1 distributions for fixed MTV values allowed us to assess the influence of water content on the R1 contrast.

### Correlations between all contrast

The acquisition of our new approach is based on a protocol typically employed for estimating R1, MTV and magnetization transfer ratio (MTR)^9,20,43,44^. Consequently, we assessed how distinct are the new qMRI parameters we propose; the tissue relaxivity, the SWF and p(R1|MTV=0), in comparison to standard qMRI parameters (R1, MTV and MTR). For this aim, we evaluated the voxel-wise correlations between parameters across the whole brain (Sup. Fig. 6). Inherently, the correlations between qMRI parameters are relatively high. For example, the correlations of R1 with MTV and MTR were substantial (R^2^=0.82 and R^2^=0.72 correspondingly), demonstrating the large similarity shared among standard qMRI parameters. Among the correlations with the new parameters, the highest correlation was observed between the tissue relaxivity and R1 (R^2^=0.8). SWF and p(R1|MTV=0) also displayed some correlations with MTR and R1, although to a lesser extent (R^2^<0.6). Notably, the weakest correlation among all tested comparisons was observed between the SWF and the non-water content estimated by MTV (R^2^=0.18). Many qMRI parameters exhibit high sensitivity to the amount of water, a factor that significantly influences the pronounced MR contrast between gray-matter and white-matter tissues^14,15,19^. By definition (eq. 19), and as evident by the low correlation of the SWF with MTV, the SWF does not depend on the absolute amount of water but rather on the fraction of saturated water, thereby highlighting distinct characteristics of the water environment within the voxel.

### White matter hyperintensities

Finally, we provide a demonstration for the applicability of the proposed qMRI technique. For this aim, we present preliminary evidence supporting the feasibility of our approach in analyzing a subject with white matter hyperintensities (WMH, see Fig. 8a for the FLAIR image). We show that our approach provides an ample of qMRI maps that allow an informative analysis of the WMH pathology. Figure 8 illustrates the various contrasts generated by our method within white matter lesions. Additionally, we conducted a comparison between measurements in the normal-appearing white matter (NAWM) and in WMH. We found a decrease in MTV and R1 within WMH compared to NAWM (Fig. 8b). This suggests tissue loss and an increase in water content within these lesions. Our approach complements these findings by offering additional biophysical information (Fig. 8c). Specifically, the SWF decreased within the lesions compared to NAWM. Therefore, while there is more water in the tissue, its saturation is less efficient. Examining the conditional distribution p(R1|MTV=0), reveals the residual R1 not explained by MTV. Interestingly, we identified a higher residual R1 in WMH compared to NAWM, implying that sources other than myelin might have a greater impact on R1 relaxation within the lesions. Furthermore, a decrease in tissue relaxivity in WMH was observed. This suggests an altered lipids and macromolecules composition which is less efficient in inducing relaxation. Notably, the largest effect size between WMH and NAWM, among all qMRI parameters tested, was found for the tissue relaxivity. This finding was replicated on five subjects with WMH (Sup. Fig 7), emphasizing the potential of the tissue relaxivity for studying white matter abnormalities.

**Figure 8:**
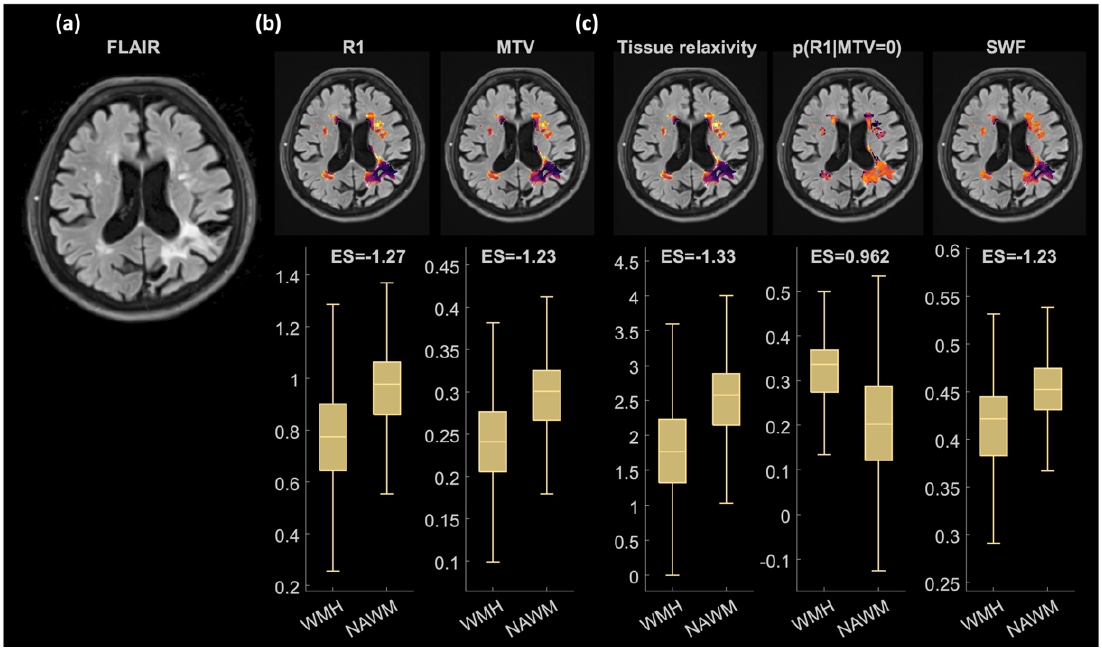
Demonstration of the new qMRI contrasts obtained in a representative older subject with WMH lesions. **(a)** The FLAIR image used for segmenting the WMH lesions in a representative subject **(b)** Differences in existing qMRI parameters (R1 and MTV) between WMH and NAWM. Upper row: the quantitative contrasts in the lesions overlaid on the FLAIR image (values were z-scored and are between [-2 2]). Lower row: boxplot of the differences between WMH and NAWM across voxels in a single subject. ES=effect size (based on Cohen’s d). **(c)** Similar analysis for the tissue relaxivity, p(R1|MTV=0) and SWF within the WMH lesions and in NAWM. These parameters supplement the existing qMRI analysis of WMH with R1 and MTV by providing additional biophysical information. Parameters were calculated by the fast implementation of our approach. For each box; the 25th, 50th, 75th percentiles and extreme data points are shown.

## Discussion

We introduce here a new approach for *in vivo* estimation of distinct water populations and tissue relaxivity within the sub-voxel environment. First, we show that this approach produces distinct contrast in the *in vivo* human brain. Consistent with our theoretical hypothesis, using MT to saturate some of the signal-producing water protons results in a lower estimation of the effective water content. Additionally, measuring less water within the voxel effectively increases the apparent tissue fraction. The MT saturation also changes the relaxation properties within the voxel, leading to an increased rate of longitudinal relaxation.

The theoretical model we use^32–34^ includes two fairly strict assumptions, that may not translate to *in vivo* data^20,46^. First, it assumes there is no direct saturation of the liquid spins. We propose a new approach to estimate and correct for direct saturation effects, based on measurements in the ventricles. This calibration significantly improves the alignment of *in vivo* results with the theory, demonstrating its importance. Notably, this calibration method could be applied to other MT-based qMRI approaches as well, highlighting the generalizability of our findings.

The second assumption of the model is that the macromolecular spins are fully saturated. However, examining *in vivo* data, we find that there is only partial saturation of the macromolecular spins. Consequently, parameters estimations vary with the strength of the MT pulse. This limitation should be considered when comparing parameters acquired with our approach for different MT pulse characteristics. Such sensitivity to the acquisition parameters and the pulse characteristics is also observed for MTR, and is expected to characterize other MT approaches as well^35,47^. This implies that other widely used MT parameters will also benefit from careful consideration of partial saturation effects^47^. Importantly, more elaborated quantitative MT techniques ^20–28,35^ allow for accurate modeling of partial saturation effects, providing a comprehensive description of the MT phenomena. These MT models often include multi-exponential relaxation of several distinct compartments, while the model we used is based on fitting a single observed exponential relaxation rate. In this study, our primary goal was to establish an MT-weighted WC mapping technique and voxel-wise tissue relaxivity estimation for the first time. Thus, our work complements previous approaches and broadens the scope of microstructural information derived from available qMRI approaches.

Despite the limitations of the simplifying assumptions in our theoretical model, we observe substantial agreement between *in vivo* results in the brain and the theoretical predictions. We also show the reproducibility of our approach in scan-rescan experiments and its robustness to changes in acquisition parameters such as TR and flip angle. Furthermore, we show that fitting of a single-exponential model to MT-weighted data is as accurate as the standard approach of fitting it to non-MT-weighted data. This suggests that the exchange between bound and free water is fast enough to produce an observed single-exponential relaxation dynamics^48^. Notably, even in the absence of MT pulse, some portion of the R1 relaxation process originates from transfer of magnetization between compartments^16,19,47,49^. Thus, any fitting of a single-exponential relaxation model to *in vivo* data should be interpreted as measuring the observed relaxation rate, which represents a combination of the relaxations and exchange rates of different compartments.

Tissue relaxivity was initially defined based on the variability in R1 and MTV across voxels, limiting its resolution to the level of whole ROIs^14^. Here, we developed an approach for estimating tissue relaxivity at the single-voxel level, producing a tissue relaxivity map for the first time. Interestingly, there is variability in tissue relaxivity between voxels within each ROI. This variability can be attributed to the increased biophysical and spatial resolution compared to the whole-ROI approach, as well as to model imperfections. Further investigation is needed to better understand the sources of this variability. However, we find strong agreement between the whole-ROI tissue relaxivity and the average voxel-wise tissue relaxivity in the ROI. Therefore, on average, similar estimates of tissue relaxivity can be obtained based on the variability in R1 and MTV across voxels or through changes in R1 and MTV with MT. This further strengthen the effectiveness of our model in providing accurate tissue relaxivity maps. Tissue relaxivity has been associated with the molecular composition of brain tissue. By measuring tissue relaxivity within the sub-voxel microstructural environment, we can obtain a more comprehensive description of the brain tissue’s lipid and macromolecular landscape.

Our technique contributes to existing MRI approaches by producing a new set of distinct parameters based on standard multi-parametric acquisition and combination of several qMRI parameters. Technically, any qMRI protocol that includes at least two variable flip angle SPGR scans and at least one MT-weighted SPGR scan, such as the widely used MPM protocol, can be used for implementing our approach. Importantly, our technique improves the biophysical interpretation of tissue characteristics in qMRI. For example, the SWF indicates the fraction of saturated water relative to total water. We show that, unlike most qMRI parameters which are highly sensitive to the overall water content in the tissue^14^, the SWF is more sensitive to the distribution of the water populations within the voxel. Additionally, we quantify these different water populations, by mapping the saturated and unsaturated water, thus advancing water content mapping techniques.

Furthermore, by mapping the linear dependency of R1 on MTV within voxels, we were able to estimate the expected R1 distribution for a fixed tissue fraction across the brain. We define these synthetic R1 contrast as the conditional distributions of R1. Similar to other works on synthetic MR contrasts^6,50–52^, this approach provides biophysically-informed modulation the R1 contrast. A considerable portion of the R1 variability across the adult brain, particularly between CSF, GM and WM, can be attributed to regional differences in the myelin content^53–58^. Higher myelin concentration leads to a higher tissue fraction and consequently higher R1 values. By calculating the conditional distributions of R1 for fixed tissue fractions, we can study regional differences in R1 contrast beyond myelin. For example, p(R1|MTV=0), reflects the residual R1 after accounting for the effect of myelin. Interestingly, this contrast shows higher R1 values in GM compared to WM, supporting its invariance to regional changes in myelin and implying a possible association with other tissue components such as iron^57–59^. Additionally, we observe a R1 contrast inversion between WM and GM with increasing MTV values. Intriguingly, such R1 contrast inversion is also observed during development, and is typically associated with myelination^60,61^. The conditional distributions of R1 also show unique cortical profiles, with distinct regional differences between cortical areas. Therefore, employing the conditional R1 contrasts to study various processes in the human brain can enrich our biophysical understanding of changes in brain tissue.

Finally, we demonstrate the applicability of our approach for imaging WMH. We find that it disentangles the effects of tissue degradation and molecular alterations on this pathology. Tissue degradation in the lesions can be estimated based on MTV, while tissue relaxivity captures the effects of molecular alterations. In addition, we observe lower SWF in the lesions compared to normal-appearing WM. Therefore, although WMH exhibit tissue degradation and increased water content, a smaller fraction of the water is saturated, suggesting potential changes in tissue organization. Moreover, the residual R1 after accounting for MTV (p(R1|MTV=0)) is higher in WMH, implying for the presence of other tissue components, such as iron, which are more prevalent within the lesions.

In conclusion, we revisited the pioneering two-site exchange model and implemented it for tissue relaxivity and water content mapping for the first time. We show that the *in vivo* MRI measurements conform well with theoretical predictions, and address the partial and direct saturations effects. These MRI measurements can be acquired with standard qMRI protocols, and are reproducible across scans. Thus, the proposed technique offers an *in vivo* characterization of the tissue relaxivity within each voxel, which is associated with the molecular composition of the brain. It also estimates the fraction of saturated water relative to the total tissue water, thereby expanding our ability to study the water content of the brain. Finally, our technique produces conditional synthetic distributions, for biophysically-informed investigation of the R1 contrast. As demonstrated for the case of white-matter hyperintensities, this approach may have great implications for the study and diagnosis of the *in vivo* human brain.

## Acknowledgments

This work was supported by the ISF grant 1169/20, awarded to A.A.M. S.F is grateful to the Azrieli Foundation for the award of an Azrieli Fellowship.

## Author contributions

S.F. and A.A.M. conceived of the presented idea. S.F. collected the data, wrote the manuscript and designed the figures. G.Y recruited the WMH subjects, funded the scans and arranged the ethical approval. L.C collected the WMH scans.

