## Supplemental information for "A qMRI approach for mapping microscopic water populations and tissue relaxivity in the *in vivo* human brain"

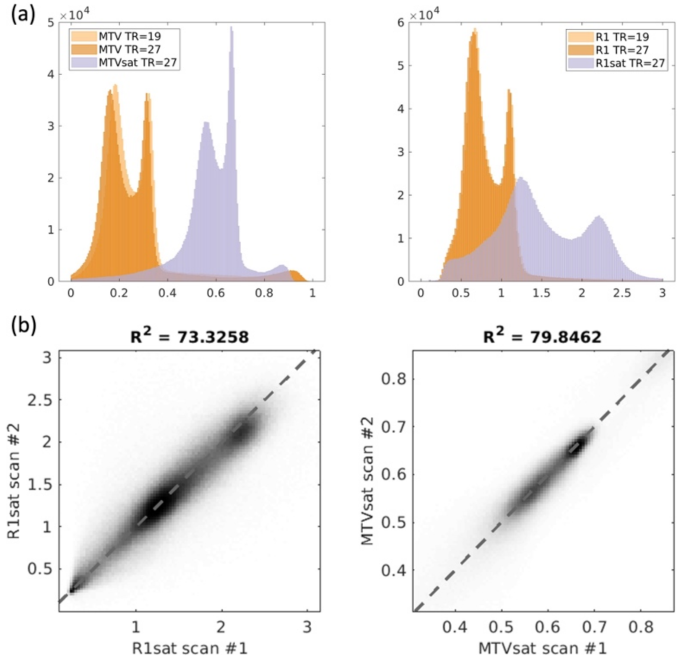


**Supplementary Figure 1: Reproducibility of our approach for different repetition times (TR) and scan-rescan experiments.** (a) MT-SPGR scans usually require longer TR than SPGR scans. We validated this is not the source of the differences between R1 to R1sat and MTV to MTVsat. Values of MTVsat (1-WCsat, left) and R1sat (right) acquired with TR=27 ms, are compared to MTV and R1 values acquired with TR=19 ms (brighter orange) and TR=27 ms (darker orange). The contrasts are shown by the distribution of values in the entire brain of a single subject. In the absence of TR difference between scans, the MT-weighted parameters are still distinct compared to non-MT-weighted parameters. Therefore, the TR difference is not the main source of the variability between contrasts demonstrated in figure 1c. (b) Scan-rescan experiment demonstrating the reproducibility of MTVsat and R1sat mapping. Data points represent all voxels in the brain of a single subject.

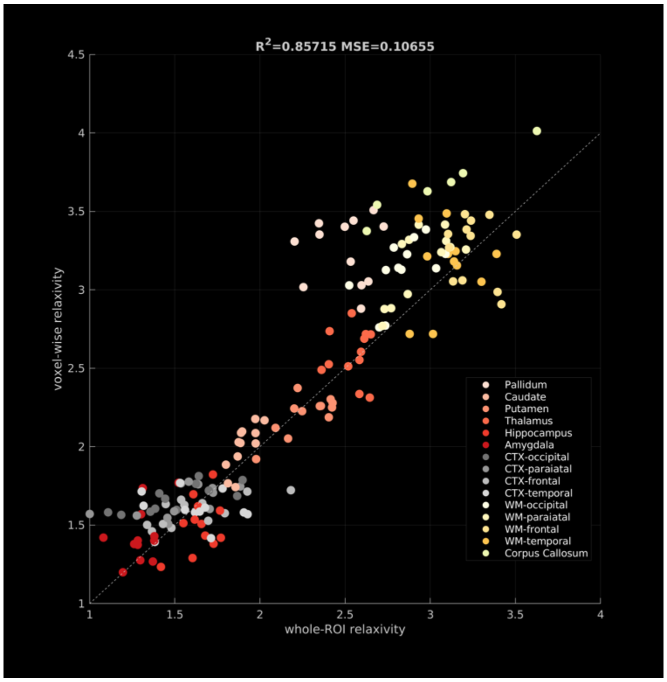


**Supplementary Figure 2: The voxel-wise qMRI map of the tissue relaxivity and its agreement with the whole-ROI approach across subjects.** The significant correlation between the averaged voxel-wise tissue relaxivity and the whole-ROI relaxivity across 15 brain regions (different colors) in 6 subjects. MSE=mean squared error.

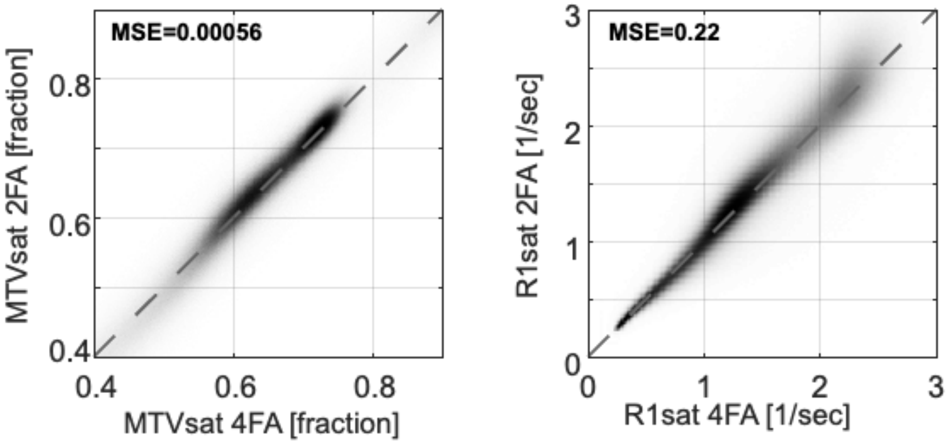


**Supplementary Figure 3: Consistency of R1sat and MTVsat mapping with varying numbers of MT-SPGR acquisitions.** R1sat and MTVsat values are derived by fitting variable flip angle (FA) MT-SPGR scans to the biophysical model (eq. 9). Parameter estimation of MTVsat (left) and R1sat (right) when using only two FAs (y-axis) is consistent with using four FAs (x-axis). Data is from all voxels in the brains of 6 subjects.

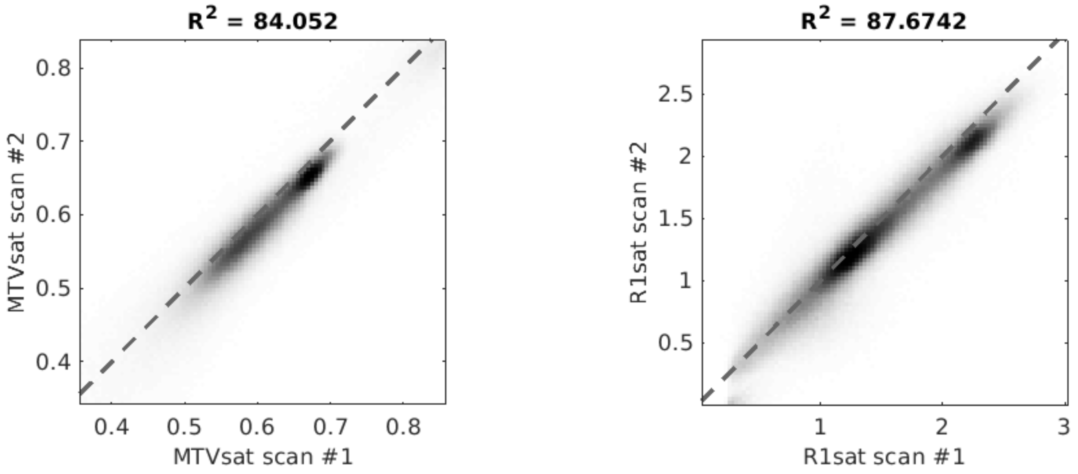


**Supplementary figure 4: The reproducibility of MTVsat and R1sat mapping with the fast implementation in scan-rescan experiments.** Scan-rescan experiment demonstrating the reproducibility of MTVsat and R1sat mapping with the fast implementation (eq. 10); MTVsat (left) and R1sat (right) maps were estimated based on variable flip angle SPGR scans, and a single additional MT-SPGR scan. A biophysical constraint reduced the number of unknown parameters (eq. 7). Data points represent all voxels in the brain of a single subject. MSE=mean squared error.

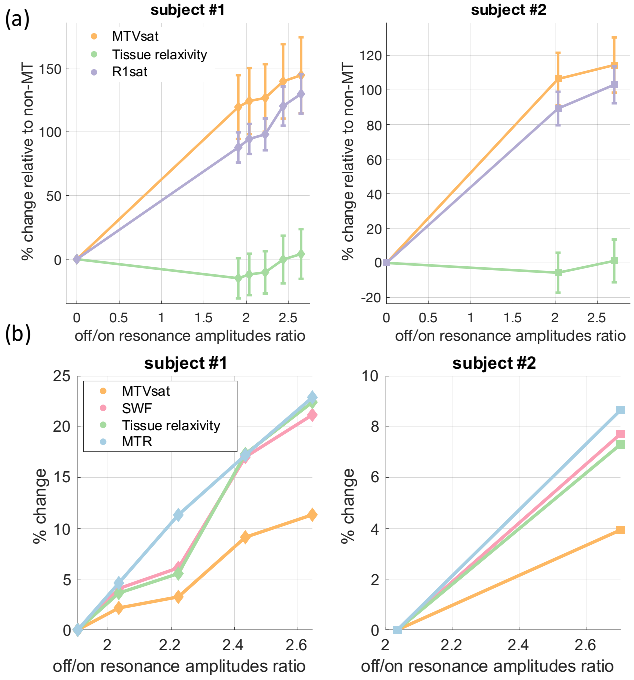


**Supplementary Figure 5: Testing partial saturation effects for two additional subjects.** (a) Comparing parameters estimated using different MT saturation strengths to those estimated without MT for two additional subjects: percentage change in R1sat, MTVsat, and tissue relaxivity (y-axis) across various MT pulse amplitudes (off-resonance amplitude to on-resonance amplitude ratio, x-axis). Changes are relative to the estimation of these parameters without MT (R1, MTV, and who;e-ROI tissue relaxivity, respectively). Parameters were calculated with the fast implementation of our approach, results are shown in the WM. Error bars indicate standard deviations. Subject #1 was scanned with a lower resolution (1.6 mm compared to 1 mm isotropic). (b) The changes in parameters estimations across different MT saturation strengths for two additional subjects. percentage change in MTVsat, voxel-wise tissue relaxivity, SWF and MTR (y-axis) for various MT pulse amplitudes (off-resonance amplitude to on-resonance amplitude ratio, x-axis). Changes are relative to the lowest MT pulse amplitude acquired. Parameters were calculated with the fast implementation of our approach, results are shown in the WM. Subject #1 was scanned with a lower resolution (1.6 mm compared to 1 mm isotropic).

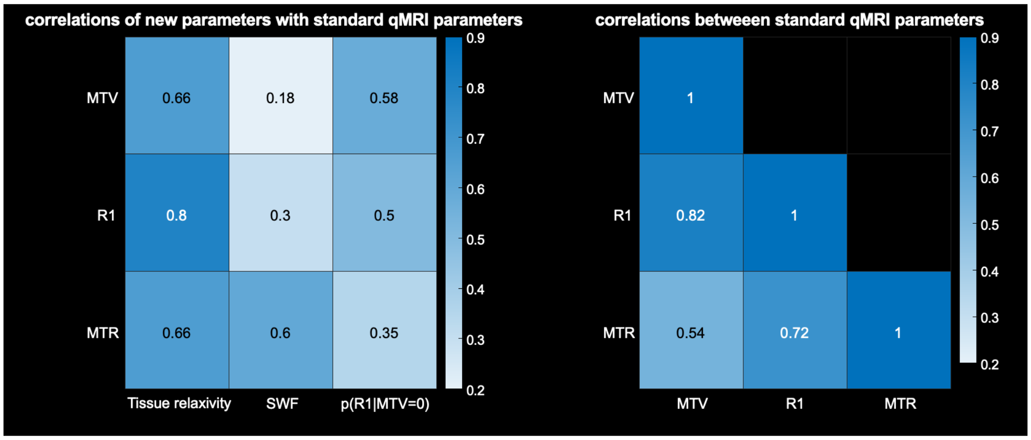


**Supplementary figure 6: The new qMRI parameters provide distinct contrasts of the human brain, different from standard qMRI parameters.** Left panel shows the correlation of the new qMRI parameters generated by our approach (tissue relaxivity, p(R1lMTV=0) and SWF) with standard qMRI parameters (R1, MTV and MTR). Correlations were computed across voxels of the whole brain, and were than averaged over n=6 subjects. Correlation matrix is color coded by the R2 of the correlations, exact values are also presented. Right panel shows similar analysis for correlations between standard qMRI parameters.

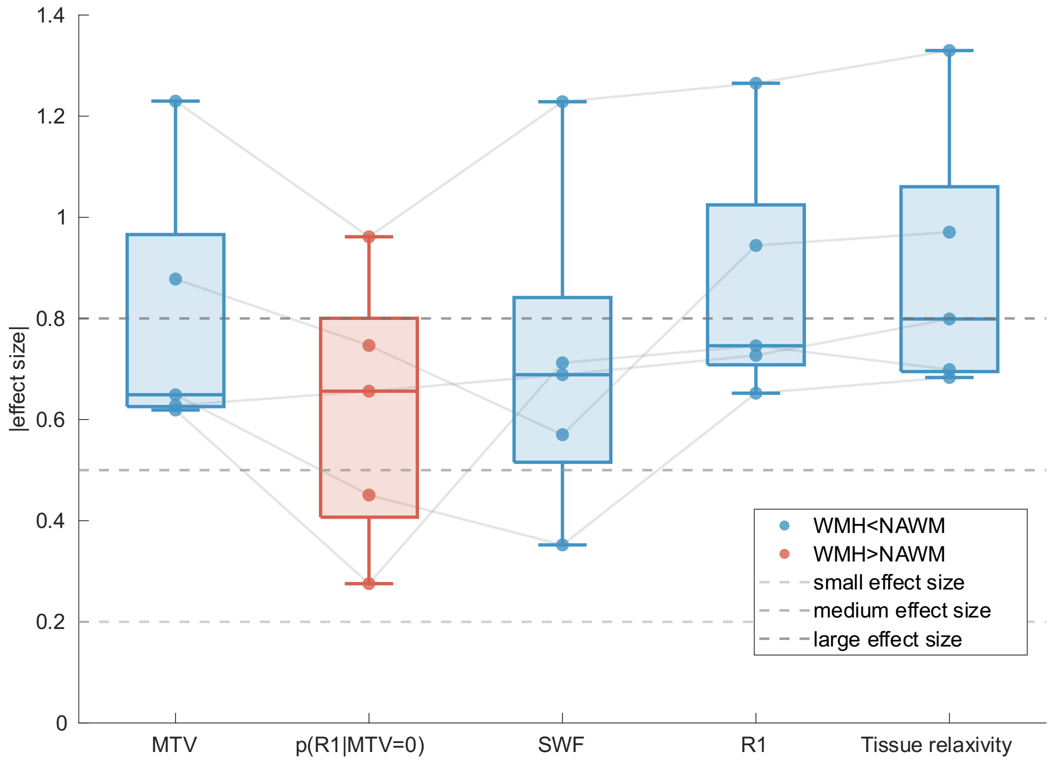


**Supplementary Figure 7: The differences between WMH and NAWM across subjects.** The effect size for comparing WMH and NAWM (Cohen's d, y-axis) for different qMRI parameters (x-axis). The effect sizes are presented in absolute values, the direction of the effect is shown by the box color. Different data points show results for n=5 subjects (lines connect between measurements of the same subject). Dashed lines indicate the significance of the effect (darker color is for larger effect). Parameters are ordered by the median absolute effect size. The largest effect size between WMH and NAWM, among all qMRI parameters tested, was found for the tissue relaxivity.

**Supplementary Movie 1: demonstration of the conditional R1 distributions.** Demonstration of the conditional R1 distributions, p(R1|MTV=x), across all possible values of MTV (x=0-1) in a representative subject. Each contrast represents the regional changes in R1 when fixing MTV values for the entire brain, and was estimated by the linear relaxivity equation (eq. 22). The dynamic range varies between frames and is adjusted between the 5th and 95th percentiles for each frame. The contrast inversion between GM and WM is observed.
